# How do plant RNA viruses overcome the negative effect of Muller’s ratchet despite strong transmission bottlenecks?

**DOI:** 10.1101/2023.08.01.550272

**Authors:** Guillaume Lafforgue, Marie Lefebvre, Thierry Michon, Santiago F. Elena

## Abstract

Muller’s ratchet refers to the irreversible accumulation of deleterious mutations in small populations, resulting in a decline in overall fitness. This phenomenon has been extensively observed in experiments involving microorganisms, including bacteriophages and yeast. While the impact of Muller’s ratchet on viruses has been largely studied in bacteriophages and animal RNA viruses, its effects on plant RNA viruses remain poorly documented. Plant RNA viruses give rise to large and diverse populations that undergo significant bottlenecks during the colonization of distant tissues or through vector-mediated horizontal transmission. In this study, we aim to investigate the role of bottleneck size, the maximum population size between consecutive bottlenecks, and the generation of genetic diversity in countering the effects of Muller’s ratchet. We observed three distinct evolutionary outcomes for tobacco etch virus under three different demographic conditions: (*i*) a decline in fitness following periodic severe bottlenecks in *Chenopodium quinoa*, (*ii*) a consistent fitness level with moderate bottlenecks in *C. quinoa*, and (*iii*) a net increase in fitness when severe bottlenecks in *C. quinoa* were alternated with large population expansions in *Nicotiana tabacum*. By fitting empirical data to an *in silico* simulation model, we found that initiating a lesion in *C. quinoa* required only 1-5 virions, and approximately 40 new virions were produced per lesion. These findings demonstrate that Muller’s ratchet can be halted not only by increasing the number of founder viruses but also by incorporating phases of exponential growth to large populations between bottlenecks. Such population expansions generate genetic diversity, serving as a buffer against, and potentially even leveraging, the effects of genetic drift.

## INTRODUCTION

Viruses have very heterogeneous genomic structures and replication strategies. Among viruses, a defining feature of RNA viruses is their high mutation rate, with estimates ranging from 10^−4^ to 10^−6^ mutations per base per replication round (1–4). During the process of evolution, natural selection can only operate on the diversity present within a population. However, beside selection, random sampling of individuals may occur during transmission events, leading to a reduction in genetic variability, which influences the evolution of a viral population (5). The strength of genetic drift depends on the effective population size (*N_e_*,) *i.e*., the abundance and genetic diversity of reproducing individuals. Small *N_e_* boosts the strength of genetic drift, magnifying stochastic effects throughout evolution (6). If virus infections are frequently initiated by a small number of virions (*i.e*., small *N_e_*), the resulting genetic bottlenecks may have a major impact on disease progression in individual hosts as well as in the evolution of virus population after multiple rounds of host infection. Furthermore, the frequency of mixed-genotype coinfections, including those by multipartite viruses, is determined by the number of infecting virions. Mixed-genotype infections can diminish pathogenesis-inducing interactions between viral genotypes while simultaneously decreasing evolutionary-relevant interactions such as recombination between distinct virus genotypes or segment segregation in segmented viruses. Although these effects are straightforward in and of themselves, their evolutionary consequences may not always be obvious (7).

The characteristics of *N_e_* are likely to affect the genetic consequences of transmissions associated with repeated bottlenecks (7). Hence, starting from a parental population in which substantial genetic diversity may exist, the sampling of a small number of individuals (founder effect) can lead to a profoundly different genetic composition in the resulting population (8, 9). If this founder effect is strong and repeated, populations can diverge rapidly, and bottlenecks tend to facilitate the random fixation of variants in populations regardless of selective value (mean fitness) (10, 11). Repeated sampling of a small number of founders take place along the infection of plants at four relevant levels. Firstly, when plants are first inoculated, infection often begins with a few viral particles. In those instances in which the virus is transmitted by insect vectors (*e.g.* aphids) the number of founders per inoculation have been experimentally quantified and ranges between 0.5 and 3.2 for the monopartite potato virus Y in pepper (12) and between 1 and 2 for the tripartite cucumber mosaic virus in tomato (13). In cases of mechanical inoculations, for example, the number of founders of the monopartite tobacco etch virus (TEV) in tobacco and pepper plants, ranged between 0.9 and 1.2 (14). Secondly, once one cells has been inoculated and replication started, the virus spreads out to neighboring cells via plasmodesmata. In this case, local transmission bottlenecks have been estimated to be in the range of 5 - 6 for the bipartite soil-borne wheat mosaic virus (15) and between 1 - 1.5 for TEV (16). Thirdly, viruses spread cell-to-cell until reaching the vascular system. From there, they follow the flux of photo-assimilates to spread out to the entire plant and induce systemic infections in new developing tissues. Additional bottlenecks have been also observed associated to these long-distance movements. For example, wheat streak mosaic virus spread from the inoculated leaf to tillers is associated with a founder effect of only 4 viral particles (17). In the same ball park, the spread of tobacco mosaic virus from inoculated leaf to systemic leaves in tobacco plants rages from 0.8 to 15.9 particles (18), with slightly higher values observed for TEV also in tobacco (5.83 to 107) (16), pea seed-borne mosaic virus (PSbMV) in pea plants (74 to 133) (19); and cauliflower mosaic virus in turnips (9.6 to 190.2) (20). Fourthly, for vertically-transmitted viruses, empirial estimates of bottleneck sizes are scarce, and the only available estimate suggests a bottleneck as narrow as 0.84 infectious particles for PSbMV in pea seeds (19).

Intracellular bottlenecking processes during cell-to-cell movement (21), combined with high mutation rates, can cause the serial fixation of deleterious mutations and a concomitant decrease in mean fitness, the so-called Muller’s ratchet effect (22). In nature, *N_e_* widely varies for viruses as a function of inoculum dose and host susceptibility (23,24). In the case of plant viruses, high mutation rate may constitute a disadvantage for RNA virus survival, as most mutations shall be deleterious (25). Muller’s ratchet has been demonstrated with bacteriophage Φ6 (26), vesicular stomatitis virus (27), foot-and-mouth disease virus (28), human immunodeficiency virus type 1 (29), and TEV (30) as well as for bacteria (31) and yeast (32). In all of these studies, in the absence of sexual reproduction, genetic mechanisms such as recombination cannot compensate for the accumulation of deleterious mutations (33, 34). In very large populations, fitness recovery can also occur through the fixation of compensatory or reversion mutations (35, 36). However, when *N_e_* is small, the fitness loss due to Muller’s ratchet is directly correlated to the fitness of the ancestral population (37, 38).

So far, all experimental studies of the Muller’s ratchet effect used protocols involving passages of viruses into cultured cells or unicellular organisms. These experimental conditions were therefore a poor representation of the complexity of natural hosts and possible vectors (39–46). Multicellular organisms display structures made of various differentiated tissues unevenly distributed in space and submitted to physical stress and possibly to immune responses, all these factors contributing to shape the genetic structure of infecting viral populations (40, 43). Among the organisms used for experimental verification of Muller’s ratchet effect, plant RNA viruses have been poorly represented.

TEV (species *Tobacco etch virus*, genus *Potyvirus*, family *Potyviridae*) possesses a positive-sense single-stranded RNA genome. TEV has been extensively studied from an evolutionary perspective (5), including its mutation and recombination rate (47, 48), mutational effects (49, 50), and epistasis (51) as well as its replication dynamics within its host (16). The virus induces local lesions on *Chenopodium quinoa* WILLD and a systemic infection in its natural host *Nicotiana tabacum* L. It has been reported that repeated transfers of local lesions generated by TEV in *C. quinoa* leads to a significant reduction of the Malthusian growth rate (30), a measure of absolute fitness. Because the virus is not moving systemically within the plant, local lesions can be used to separate a small subpopulation of viruses from the original pool (54): another characteristic of this model is a putative moderate virus multiplication rate due to the plant hypersensitive response that results in the necrotic lesions (55). Factors governing evolution in complex multicellular hosts systems are multiple. However, by the use of two host (*N. tabacum* and *C. quinoa)* our experimental system allows dissecting the influence of two demographic parameters: the strength of individual bottlenecks and the amount of replication events between two consecutive bottlenecks, in a complex multi-cellular host plant. The literature highlights the influence of fluctuating environment on mutations (56, 57). In this experiment, *C. quinoa* serves as a valuable tool for isolating and manipulating infectious particles. We hypothesize that the limited replication within the lesions does not promote the generation of new mutations within this particular context. Notably, the influence of altering the host is deemed negligible in this specific case, as no adaptation (systemic infection) has been observed in *C. quinoa*, leading to assume that observations in this study is principally lead by the effective population size combined to multiplication of the virus.

In this study, capitalizing on a previous study using the TEV/*C. quinoa* pathosystem (30), we investigated the impact of these two demographic parameters on this potyvirus infectivity in its natural host (*N. tabacum*), and how these factors impact the recovery or decrease of infectivity in the viral progeny.

## MATERIALS AND METHODS

### *In vitro* RNA transcription and plant inoculation

The pTEV-7DA infectious clone, provided by Prof. James C. Carrington (Donald Danforth Plant Sciences Center, St. Louis MO, USA), was the source for the ancestral virus for all experiments (45). Infectious plasmids were linearized with *Bgl*II (Fermentas, Vilnius, Lithuania) and transcribed into 5’-capped RNAs using SP6 mMessage mMachine^®^ Kit (Ambion Inc., Austin TX, USA). Transcripts were precipitated (1.5 volumes of DEPC-treated water, 1.5 volumes of 7.5 M LiCl, 50 mM EDTA) and resuspended in DEPC-treated water. RNA integrity was assessed by gel electrophoresis and its concentration quantified spectrophotometrically with a NanoDrop (Thermo Fisher Scientific, Waltham MA, USA). The obtained RNA was used to inoculate groups of 4-week-old *N. tabacum* cv. *Xanthi* plants by means of abrasion of the third true leaf with 4 μg of RNA (Fig. 1). Plants were maintained in the green house at 25 °C and 16 h light for one week. Symptoms appeared 5 - 6 days post-inoculation (dpi).

**Fig. 1.**
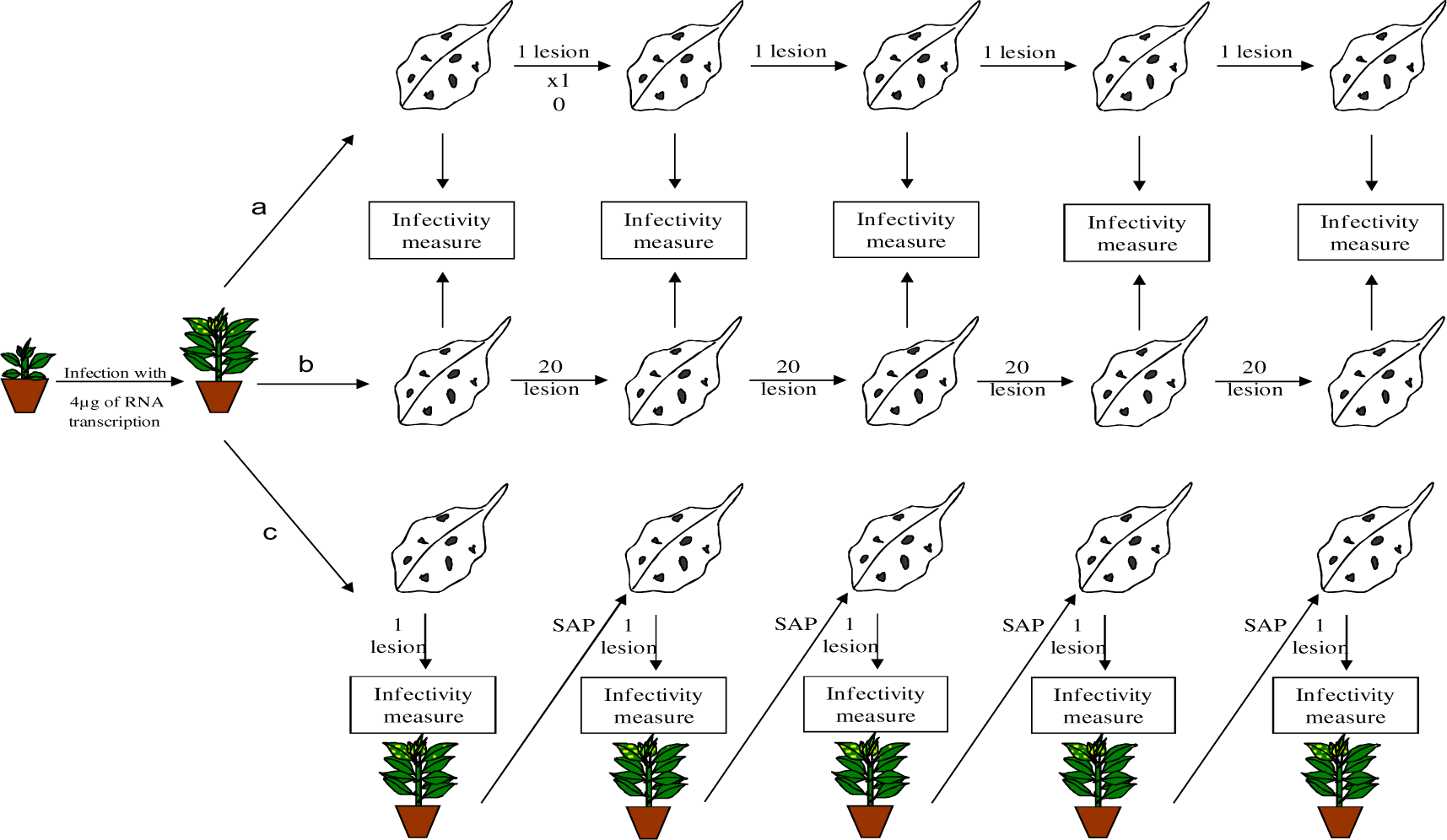
Schematic representation of the serial transfer of a TEV population derived from an initial clone. At each transfer, either a random lesion (a, strong bottleneck, ‘smallest *N_e_*’ treatment) or twenty lesions (b, weak bottleneck, ‘intermediate *N*_e_’ treatment) were used to continue serial passages. In the third ‘large *N_e_*’ treatment (c) one lesion was used to initiate infection in *N. tabacum* and then transferred on *C. quinoa*. After each transfer, the population fitness was measured on *N. tabacum*. Each evolutionary treatment was replicated 10 times.

### Inoculation of *C. quinoa* and generation of mutation-accumulation lineages

An initial sap was prepared with 1 g of infected tissue (from *N. tabacum* plants 7 dpi) ground in 1.5 mL of phosphate buffer (50 mM phosphate, 3% polyethylene glycol 8000). Four different leaves from five different 4-week-old *C. quinoa* plants were rub-inoculated with 10 μL of sap. Lesions on inoculated leaves were harvested at 10 dpi. A well-isolated lesion was perforated from the leaf using the wide end of a 5 μL micropipette tip (approximately 4 mm diameter disc) and ground in a mortar with 24 μL of phosphate buffer. The extract was carefully collected, centrifuged for 1 min at 12,000 g and the supernatant was used for inoculation.

The experimental manipulation of the bottleneck size was done using two different modalities (Fig. 1). The ‘smallest *N_e_*’ treatment was used to apply a strong bottleneck. Briefly, 10 lesions were processed separately to create ten independent single lesion lineages: 5 μL of each lesion sap used to inoculate four different leaves of *C. quinoa* to generate the next generation of lesions. After 10 dpi, one lesion was used to generate the next generation. To generate a weaker bottleneck effect, the ‘intermediate *N_e_*’ treatment was used. Ten independent pools of 20 lesions were pooled and grounded together in 240 μL of extraction buffer producing ten independent group lesion lineages. The same procedure was used as for single lesion lineages (*i.e*., 10 μL of lesion sap were used to inoculate ten *C. quinoa* leaves).

To introduce a larger number of generations between consecutive bottlenecks, the ‘large *N_e_*’ treatment was used. Briefly, in between passages in *C quinoa*, the progeny was sequentially inoculated in TEV natural host, *N. tabacum*. After 10 dpi, a *N. tabacum* infected plant was ground in phosphate buffer (0.5 g in 500 µL) and used to inoculate ten *C. quinoa* leaves (10 µL).

It has been previously reported that single lesion lineages tend to have a significant loss of absolute fitness until, in some cases, the lineages become extinct (30). After five passages, the experimental study ended with a loss of four lineages and a production of less than five lesions.

For all experimental treatments, lesions sampled at each passage were stored at −80 °C to perform all infectivity test later into a single block. No effect of storage at −80 °C on single lesion infectivity was observed (data not shown).

### Infectivity measurement

The fitness of each evolved lineage within each history (small, intermediate or large *N_e_* treatments), was assessed by infectivity measurement at each time point (*i.e*., the end of a given passage). The analysis of infectivity was performed on *N. tabacum*. From a given lesion on *C. quinoa*, only two tobacco plants can be inoculated. Consequently, to be statistically relevant, for each passage, up to 10 single lesions were randomly selected for each lineage. The lesions were ground in 24 µL of phosphate buffer, and 10 µL were used to inoculate the third true leaf of two *N. tabacum* plants, for each lesion. Presence of symptoms was recorded at 9 dpi. Hence, per history (small, intermediate or large *N_e_* treatments), a total of 200 tobacco plants were inoculated per passage. Infectivity was expressed as the frequency of infected symptomatic plants out of the number of inoculated ones.

### Statistical analyses

The linear regression of infectivity in *N. tabacum versus* passage number, fixing the origin at the 0.261 ±0.035 (±1 SEM) infectivity value measured at the first passage (see below), returns a slope that represents an estimate of the rate of infectivity evolution. These rates of evolution were determined for each lineage of the small, intermediate and large *N_e_*. A one-way robust Welch ANOVA was used to evaluate the effect of demographic conditions in the rates of infectivity evolution. The magnitude of the effect was evaluated using the *η*^2^ statistic; usually *η*^2^ > 0.15 are considered as large effects. Variance components were estimated by maximum-likelihood. These statistical tests were conducted with SPSS version 28.0.1.1 (IBM Corp., Armonk NY, USA).

### An *in silico* model of the growth processes taking place within single lesions

To evaluate the demographic processes occurring within a local lesion during virus multiplication, and to mimic the mechanisms acting for the small or intermediate *N_e_* treatments, we implemented a simulation model in R version 3.6.1 (https://doi.org/10.5281/zenodo.8199726). Briefly, the experiment was simulated in three steps: (*i*) generation of lesions in *C. quinoa* from infected *N tabacum*, (*ii*) multiplication within a random lesion and (*iii*) generation of lesions either from a single (small *N_e_* treatment) or from 20 digital lesions (intermediate *N_e_*treatment). Steps (*ii*) and (*iii*) were repeated four times with 200 replicates for each model.

These models take into account the following postulates: (*i*) a few number of particles shall be enough to initiate a lesion (14), (*ii*) two types of viral particles are considered: type A virions which can reproduce either in *C. quinoa* and *N. tabacum* and type D virions which can reproduce only in *C. quinoa* but had lost their capacity to infect the natural host, and (*iii*) the percentage of virions A in a lesion determines the probability of infection in *N. tabacum* defined as the infectivity rate:

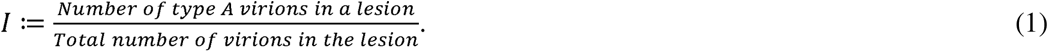

Result for each passage is given by the mean of *I* for 12 random lesions.

The parameters of the model are the number of virions able to initiate a lesion *V* ∈ (*V_min_*, *V_max_* | *V_min_*< *V_max_*) that take integer values in the range [0, 8] and the rates of replication in *C. quinoa* of type A virons 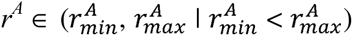 and 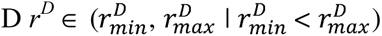 that take integer values in the interval [1, 50].

The small *N_e_* model was implemented as follows. First, a random number of founders *V* is determined. Then, among these founders, the number of virions of types A and D are derived from equation (1), *I* being given as the infectivity measured experimentally at the first passage (*i.e*., 0.261 ±0.035; see below). Replication rates in *C. quinoa r^A^* and *r^D^* are used to calculate the next pool of virions (*G* + 1) able to generate the next lesion. Twelve simulated lesions are generated. A random number in the interval (*V_min_*, *V_max_*) of founder particles is attributed to each lesion. From the *G* + 1 pool, the subpopulations of particles types A and D within each lesion was randomly assigned. For this model, a single lesion was randomly picked to repeat the process until the fourth passage. An exhaustive search in the parameter space values 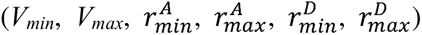 was done, resulting in 5.8·10^7^ different combinations in the case of the small *N_e_*model. For each of these simulations, 200 replicate runs were performed. Combinations of parameters resulting in estimates of infectivity and intra-lineages genetic component of variance within the observed experimental 95% confidence intervals were retained for further analyses. Furthermore, simulated rates of infectivity evolution for these parameter combinations were obtained and contrasted against the 95% confidence intervals empirically estimated. This step results in 1.8×10^7^ combinations corresponding to measured values in the small *N_e_* treatment.

The intermediate *N_e_* model uses the same canvas as for the small *N_e_* model described above and incorporates the 1.8×10^7^ combinations previously retained. The difference resides in the generation of 20 lesions subsequently pooled together, the resulting mix being used to generate the *G* + 1.

Due to the objective of the simulations, which is to monitor the parameters associated with the virus’ life cycle within a lesion, and considering the significant increase in the number of additional parameters involved in the large *N_e_* treatment (such as mutation rate and multiplication within tobacco), it was not considered in the analysis.

## RESULTS

### Critical infectivity can be restored

In all experiments, fitness was considered to be directly related to the capacity of the virus to initiate disease in *N. tabacum*. Inoculation of *N. tabacum* with sap from infected tissue of *N. tabacum* results in nearly 100% infection of plants inoculated (data not shown). After a single passage of TEV in *C. quinoa,* infectivity in *N. tabacum* was reduced to 0.261 ±0.035. This resulting virus population was then used for subsequent evolution experiments.

Serial passages of single lesions (small *N_e_* treatment) produced in *C. quinoa* resulted in a further decrease in infectivity in *N. tabacum* (Fig. 2A). Indeed, the average infectivity among lineages after five passages was 0.174 ±0.078, representing an average net reduction of 33.5% relative to the first passage. After removing four outlier values, the average rate of infectivity evolution among these lineages was −0.033 ± 0.005 per passage, which was found to be significantly negative (*t*_17_ = −7.390, *P* < 0.001). Without removing outliers, the rate of infectivity evolution was −0.018 ±0.009 per passage, indicated that it was still significantly negative (*t*_21_ = −2.034, one-tailed *P* = 0.027).

**Fig. 2.**
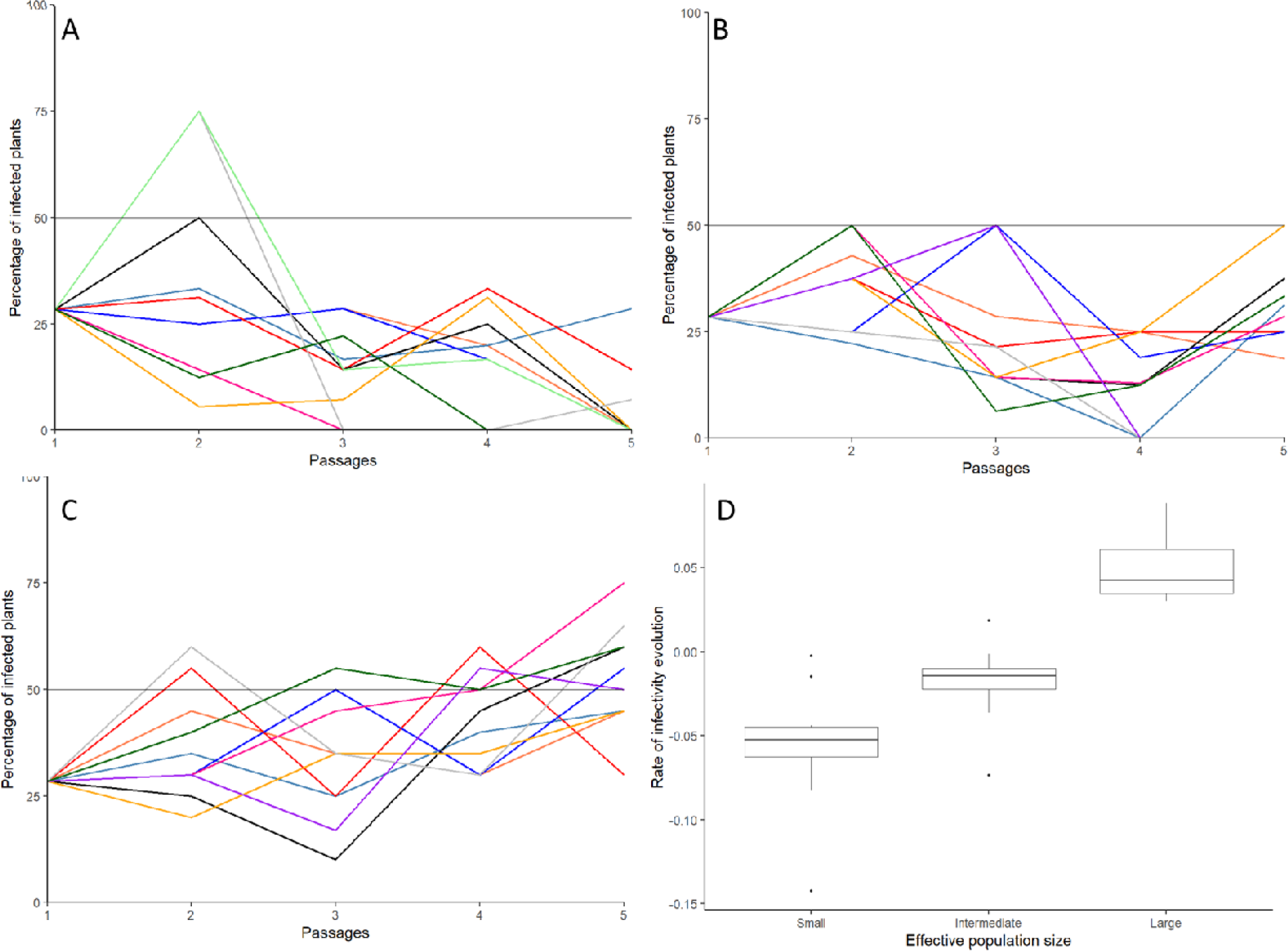
(A) Evolution of average infectivity among independent lineages at each bottleneck event (passage) for small *N_e_* = 1 (lesion-to-lesion), (B) intermediate *N_e_* = 20 (20 lesions-to-20 lesions) and (C) large *N_e_*(population expansion in *N. tabacum*) experimental treatments. (D) Box plot for the estimated rates of infectivity evolution. Error bars represent ±1 SEM.

When passaging was performed with pools of 20 local lesions (intermediate *N_e_* treatment; Fig. 2B), the final average infectivity among the 20 lineages was 0.200 ±0.076, which represents a further reduction of 21.6% in infectivity relative to the first passage. For this treatment, the rate of infectivity evolution was −0.017 ±0.006 per passage, a value that is also significantly negative (*t*_18_ = −2.654, *P* = 0.008). Finally, when passages included a population expansion in *N. tabacum* and the bottleneck in *C. quinoa* (large *N_e_* treatment), a net increase of infectivity over passages was observed (Fig. 2C), with an average final infectivity of 0.447 ±0.038 and an average rate of infectivity evolution estimated to be 0.037 ±0.005 per passage, a value significantly positive (*t*_9_ = 7.406, *P* < 0.001).

Significant differences among the three treatments exist in the rates of infectivity evolution (Fig. 3D: *F*_2,31.281_ = 28.122, *P* < 0.001), being them of very large magnitude (*η*^2^ = 0.321). However, these differences were mostly driven by the differences between the large *N_e_* treatment and the other two (Bonferroni *post hoc* test: *P* < 0.001 in both comparison), but not among the small and intermediate *N_e_*treatments (Bonferroni *post hoc* test: *P* = 1.000).

**Fig. 3.**
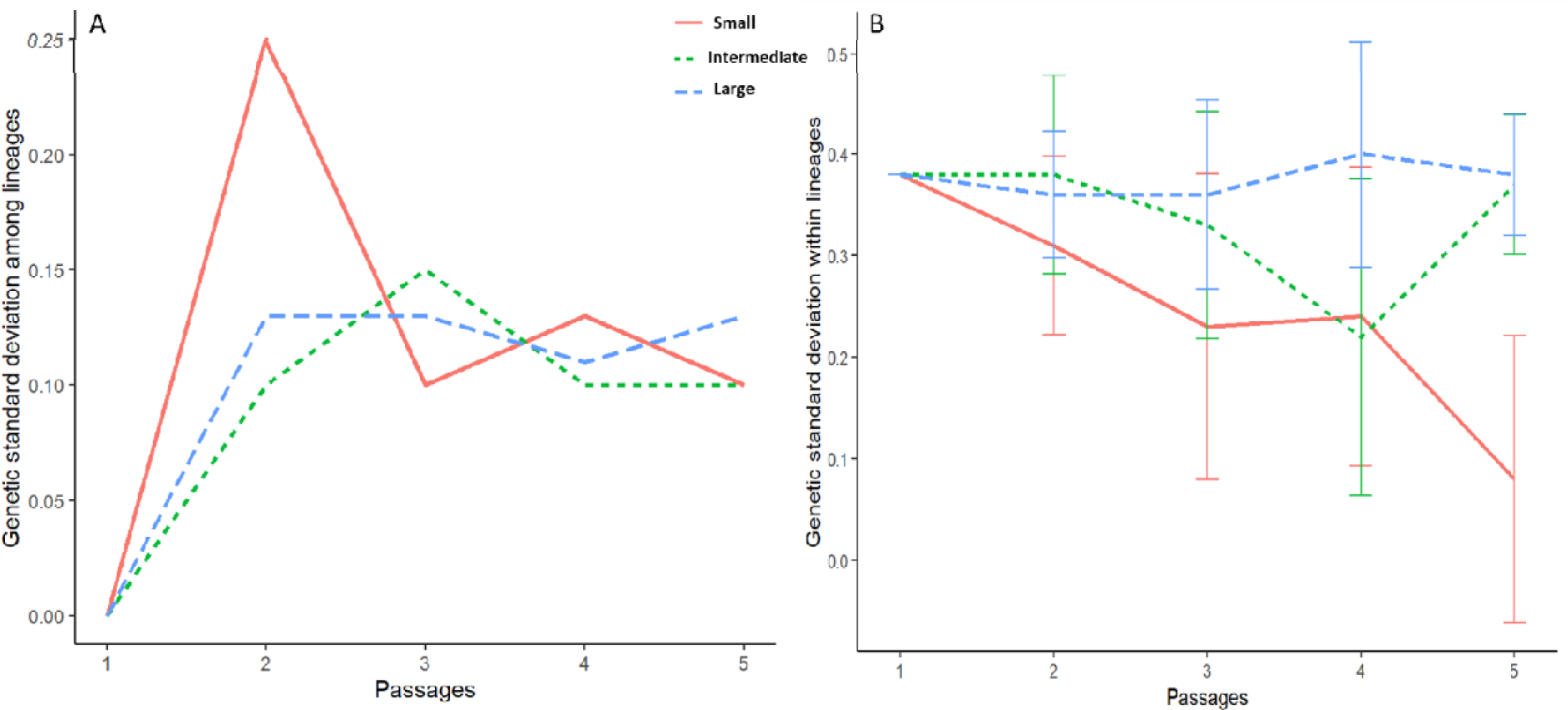
Evolution of the genetic standard deviation among (A) or within (B) lineages along the passages.

### Variance intra- and inter-lineages

For a given demographic setup, the lineage fitness can vary in *N. tabacum* due to several factors (adaptation, chance, contingency…). To evaluate the contribution of true genetic differences to the evolution of infectivity, we decomposed the total observed variance for lineages evolved in the same conditions into their additive (among lineages) and random (within lineages) components. Fig. 3A shows the evolution of the additive (*i.e*., genetic) standard deviation for the three experimental treatments. In all three treatments, genetic differences among lineages increased after the first passage, inversely proportional to *N_e_*: random fixation of alleles associated with strong bottlenecks contributes to inflating the genetic component of variance. However, throughout the experiment, subsequent strong bottlenecks resulted in a loss of genetic differences, stabilizing the variance.

Interestingly, at the beginning of the experiment, the intra-lineage variation among the lineages with the smallest *N_e_*appeared to be higher compared to the lineages derived from the other two modalities. However, there is no differences between the smallest *N_e_* and the two others modalities (Wilcoxon signed-ranks test: *W* = 7.5, *P* = 0.346 and *W* = 14.5, *P* = 0.753, respectively). At the third passage, the inter-lineage variance of the smallest *N_e_* lineages decreased and reached a similar level as the inter-lineage variance observed in the intermediate and large *N_e_* lineages. Indeed, no statistically significant effect of passages within the three different modalities has been found (Friedman test: χ^2^ = 8.8, 4 d.f., *P* = 0.066). This suggests a convergence in terms of genetic diversity among the different lineages, regardless of their initial *N_e_* values.

The analysis of lineages with varying *N_e_* revealed interesting trends in intra-lineage variation over the course of passages. Specifically, the lineages with the smallest *N_e_* exhibited a significant decreasing slope of −0.0675 in intra-lineage variation across passages (Fig. 3B: *t*_38_ = −3.870, *P* < 0.001). This finding suggests that there was a reduction in genetic diversity within those lineages over time as they were serially passaged. On the other hand, the intermediate and large *N_e_* lineages did not display any significant changes in intra-lineage variation throughout the passages (*t*_39_ = 1.039, *P* = 0.305 and *t*_37_ = −1.004, *P* = 0.322). The lack of significant variation in these lineages suggests that their genetic diversity remained relatively stable over time. This could be attributed to their larger *N_e_*, which may have provided a buffer against genetic drift and enabled the preservation of diverse genetic traits within these lineages.

### In the small *N_e_* treatment, infectivity is independent from multiplication in *C. quinoa*

The observed decrease in infectivity in *N. tabacum* after serial passages of single lesions from *C. quinoa* (small *N_e_* treatment) may be attributed to factors related to the quantity and infectivity of virions present in the inoculum derived from *C. quinoa* lesions.

It has been established that the number of lesions in *C. quinoa* is closely associated with the viral load (30), which refers to the quantity of viral particles present within the infected host. Therefore, in the context of the small *N_e_* treatment, the decrease in infectivity observed in *N. tabacum* could be linked to a lower quantity or even an absence of virions in the inoculum derived from *C. quinoa* lesions, which are capable of infecting *N. tabacum*. However, upon further investigation, no significant correlation was found between the number of lesions and the infectivity in *N. tabacum* (Spearman’s rank correlation coefficient *r_S_* = −0.017, 118 d.f., *P* = 0.854). This finding indicates that the infectivity rate in *N. tabacum* is influenced more by the genetic characteristics of the virions rather than the concentration of the viral load (Fig. 4). In other words, the genetic properties of the viral particles present in the inoculum may play a more critical role in determining the infectivity in *N. tabacum* than the sheer quantity of the viral particles.

**Fig. 4.**
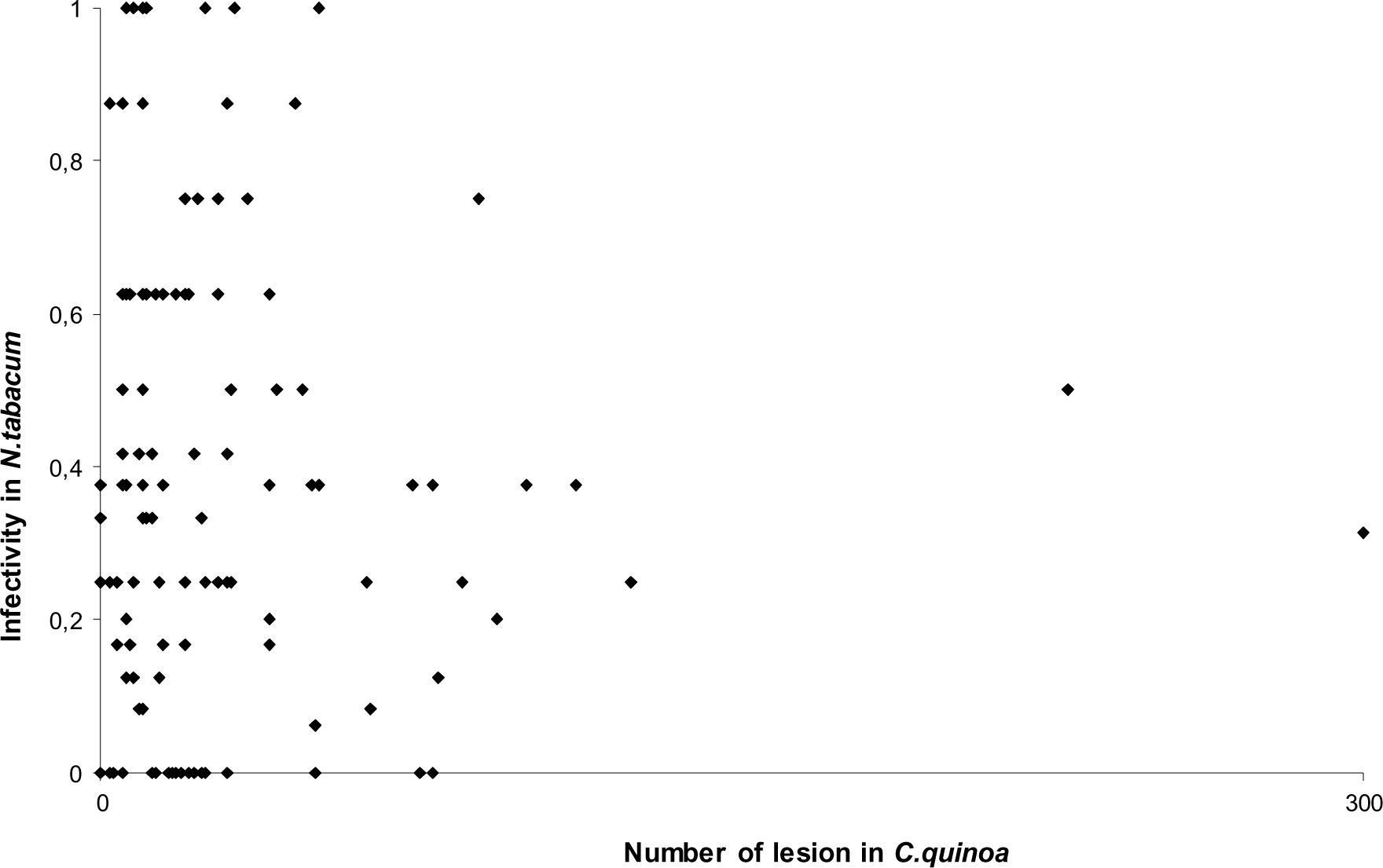
Relationship between infectivity on *N. tabacum* and number of lesion on *C. quinoa*.

### Evaluation of growth parameters for individual lesions

From the 4.2×10^7^ different combinations of parameters tested, 1.74×10^3^ (*i.e*., 0.04%) generated mean infectivity values and non-additive standard deviations within lineages (*i.e*., true noise) lying within the observed 95% confidence intervals, thus being compatible with the experimental observations for the small and intermediate *N_e_* treatments.

Fig. 5 summarizes the results of the exhaustive search of space parameters compatible with the observed results. The modes of these six distributions can be taken as the most representative values for the model parameters: *V_min_* = 1, *V_max_*= 4, = 11, = 37, = 21, and ≥ 50. These parameters show that very few particles are enough to generate a necrotic lesion in *C. quinoa* and that the D mutant types show higher replication rates in *C. quinoa* than the A types, with a replicative advantage that varies between 1.35- and 1.91-fold. To qualitatively confirm the agreement between observed data and the estimated parameter values, we run 20 independent simulations with the parameters above (Fig. 6).

**Fig. 5.**
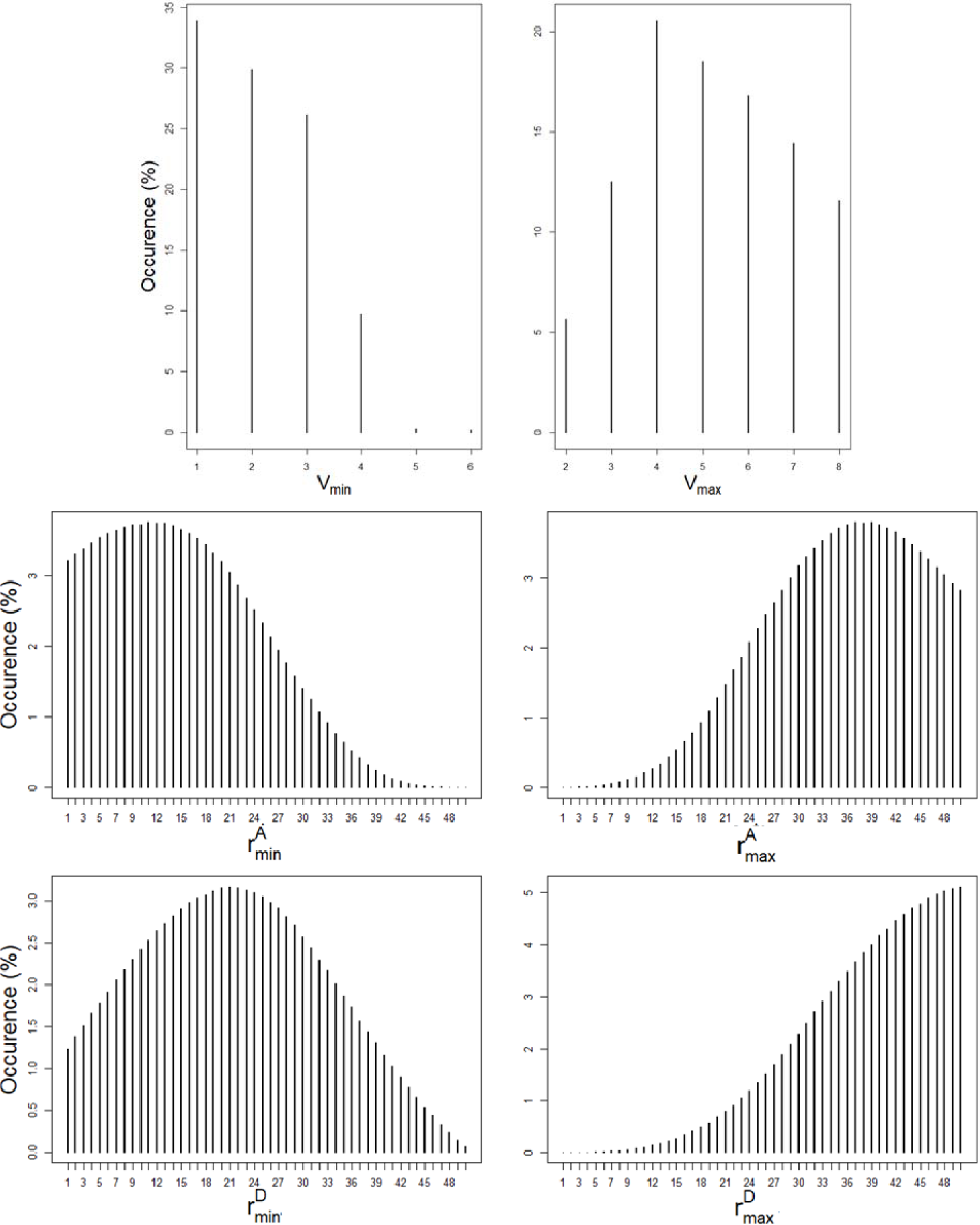
Distribution of parameters (*V_min_*, *V_max_*, , , ,) compatible with the experimental observations made for small and intermediate *N_e_* demographic regimes.

**Fig. 6.**
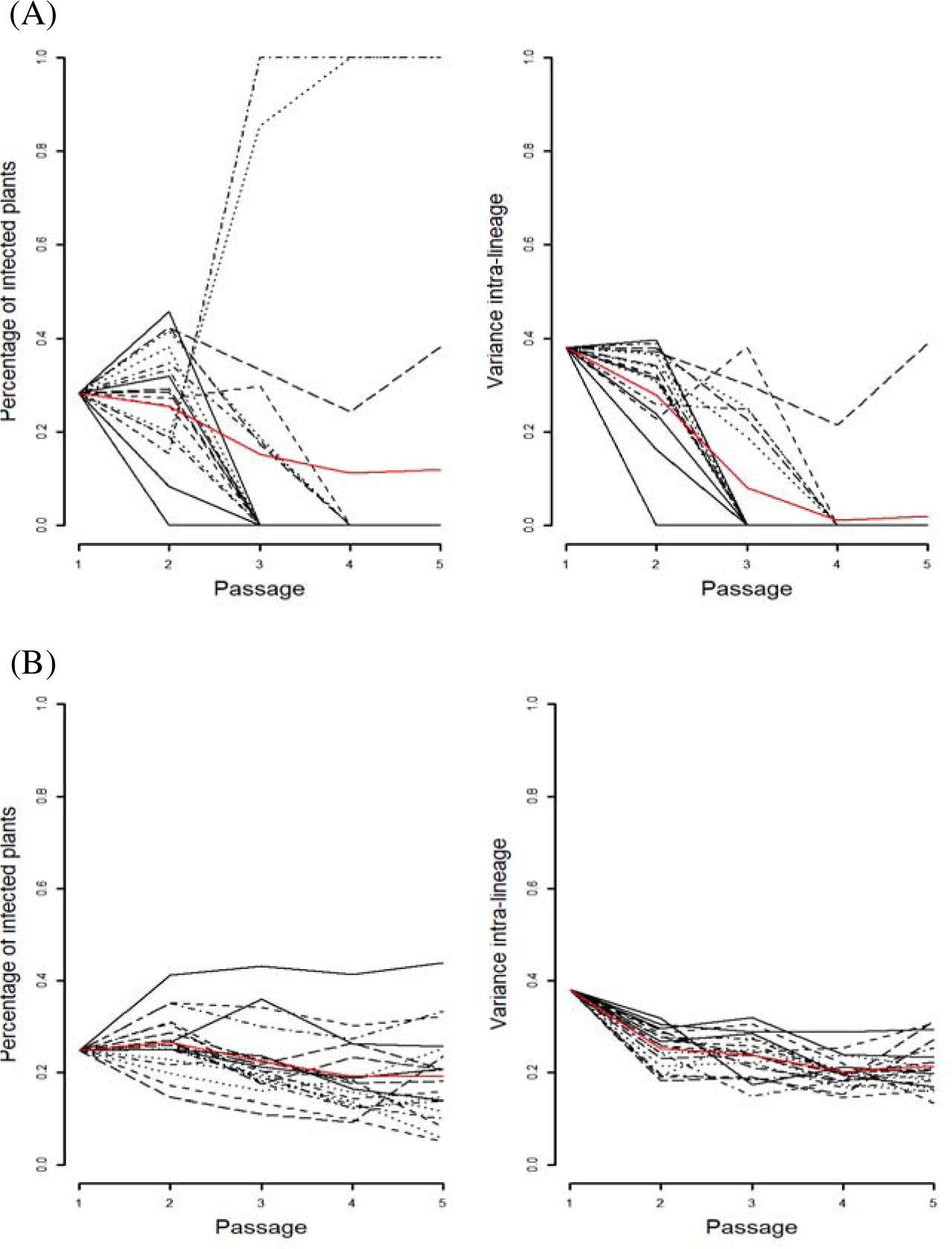
Twenty simulated lineages with the most common set of parameters. The infectivity and within-lineage error are plotted for (A) small *N*_e_ and (B) intermediate *N_e_* treatments. Red lines represent the mean of the 20 replicates.

The simulation results shown in Fig. 6A are in good agreement with the observed dynamics shown in Fig. 2A and Fig. 3A: an overall trend of infectivity to decline along passages and a strong decrease in intra-lineage variance, respectively. However, some differences exist between experimental and simulation results. Firstly, most of the simulated lineages lost their infectivity after the third passage while the majority of experimental lineages were still infectious in *N. tabacum* after the fifth passage. Secondly, some simulated lineages reached 100% infectivity in *N. tabacum*, something that was not observed in our experiments. Regarding the simulations for the intermediate *N_e_* treatment (Fig. 6B), the 20 simulated lineages show a minor reduction in infectivity, also in good agreement with the data shown in Fig. 2B. In this case, a qualitative difference exists between observed and simulated data. In this case, some experimental lineages reached 0% infectivity in *N. tabacum*, while the simulations never showed a lack of infectivity.

## DISCUSSION

Although Muller’s ratchet has been studied for several microorganisms, including viruses, so far, it has been experimentally addressed only for one plant virus (30). Bottlenecks such as vector-mediated transmission (12, 13, 41), cell to cell movement (15, 16) and new leaves systemic colonization (16, 17, 41, 42, 44) are common during the multiple stages of the infectious cycle of plant viruses. There is experimental evidence that the number of founders (*i.e*., viral genomes triggering infection) depends on the stage of the virus cycle affected by the bottlenecks. For instance, it was reported that from one to six viral particles infect a cell (the multiplicity of infection, MOI) (15, 16). Considering within-plant bottlenecks, the lowest *N_e_* reported was ∼ 4 founders for wheat streak mosaic virus and the largest ∼133 for pea seed-borne mosaic virus (19). It has also been shown that within-host bottleneck size can largely vary in the range 6 - 10^7^ among sink leaves (18, 20). These different bottlenecks size shape drastically the virus population. This means that the whole diversity present in a population can be lost by a single infection event, or be highly reduced for cell-cell or systemic movement. Studying how Muller’s ratchet operates on a plant virus population during its complete cycle (inoculation, movement and transmission) is therefore highly relevant to understanding its evolutionary dynamics.

In our study, each single lesion in *C. quinoa* represents a sample from a genetically complex viral population. This feature allowed us to investigate the contribution of different factors influencing the viral population. Using this approach, the role of bottleneck strength and multiplication rate on evolution could be investigated demonstrating hereby that different combinations of these two factors lead to global extinction (small bottleneck, low demographic dynamics), stabilization (intermediate bottleneck, low demographic dynamics) or rescue of a population (small bottleneck, high demographic dynamics). The generation of diversity is dependent on the occurrence of random mutations, and the total number of mutations expected to occur will depend on the number of generations (*i.e*., rounds of replication). Our experimental results implicate three general phenomena, genetic drift, generation of diversity and population size. Firstly, in the small *N_e_* situation, only a few genomes are randomly sampled every passage and then multiply at a low rate. This low multiplication in a local lesion generates low diversity. The hypothesis is thus that the principal force acting in this case is genetic drift through the Muller’s ratchet process whereby in a defined population only some individuals participate to the next generation.

Secondly, passages following the intermediate *N_e_* conditions (weaker bottleneck) maintained the original diversity generated in *N. tabacum*, but without any gain in fitness. In this modality, the missing force is the generation of diversity through a high multiplication rate and selection for efficient replication in *N. tabacum*. Regarding the results from the *in silico* experiments, we can extract a different general range of maximum multiplication rates of virions inside a lesion between the assumed virus particles A and D. This result could be explained by a better adaptation to *C. quinoa* of the defective particle (virion D). Finally, a large range of values for the replication rate in *C. quinoa* either for A or D virions can be explained by the variation due to the host’s response and its ability to generate the necrotic local lesions (55). Thirdly, these elements can be introduced by alternating passages in *C. quinoa* and *N. tabacum* (large *N_e_* treatment): despite the narrow bottleneck imposed by passaging only a single *C. quinoa* local lesion, diversity was generated rapidly and selection acted efficiently in *N. tabacum.* Such a net effect not only attenuated Muller’s ratchet effect but also induced an increase in fitness.

Muller’s ratchet has received quite a lot of attention from modelers. In his pioneering work, Haigh (34) found that the most relevant parameter to predict the speed of the ratchet was *n*_0_ = *N_e_e^−US^*, where *U* stands for the genomic deleterious mutation rate per generation and *s* for the average effect of a random deleterious mutation. If *n*_0_ ≥ 25, then mutations accumulate slowly but if *n*_0_ < 1 then the rate of accumulation is high and the frequency of the less mutated genotypic class disappears after each click. As deleterious mutations accumulate and reduce fitness, one expects that the number of reproducing individuals also decreases, further reducing the census population size, enhancing the effect of Muller’s ratchet and leading to an extinction process known as mutational meltdown (58). Later on, Kondrashov (58) showed that these conclusions were very sensitive to the assumption of independent mutational effects and showed that if mutations interact in a synergistic manner to determine fitness, the ratchet slowed down its speed; it can be halted with strong enough epistasis, allowing small asexual populations for survive without entering in the mutational meltdown regime. More recently, Bachtrog and Gordo (59) explored the role of beneficial mutations in the speed of Muller’s ratchet. If Muller’s ratchet is already clicking, mutations that might be neutral in the wild-type genotype can become fixed if they arise in a nonoptimal genotype as, since the loss of the least-mutated class implies that a suboptimal genotype can now become selectively advantageous. Indeed, these authors showed that as far beneficial mutations are rare, the rate of fitness decline is weakly affected. Here, we have modeled the experiments to explore some features of the initiation and replication inside a local lesion and even if results of simulated and experimental data are close, discrepancies could be explained by different factors. First, we capped the maximum replication rates for the viruses inside the lesion to 50. This is an arbitrary choice and this value can be larger, especially for the D virions, where the maximum seems not be attained yet (Fig. 4). Anyway, the curves of occurrence of the values of 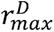 show a similar shape than observed for 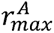, suggesting 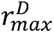 shall be close to its maximum value. In the simulation of the small *N_e_* model, two replicates show an intriguing pattern of increase in infectivity in *N. tabacum* (Fig. 5A). This suggests that several non-mutually exclusive explanations: (*i*) our plant experiments involved a limited number of lineages and thence we might have missed these exceptional cases. (*ii*) The assumption of only two types of virions (A and D) is far too simplistic and a gradient of fitness effects could exist. And (*iii*) some factors possibly implicated in the virus life cycle inside a lesion have not been well incorporated in the model, as for example the phenomenon of cooperation. Despite these differences, our model sets specific demographic parameters inside a lesion within a certain order of magnitude, which allows us to understand the virus cycle in such a particular environment. One of the first result of the model indicates that a single virion is enough to generate a lesion as previously described to initiate an infection (7, 61). The maximum number of particles initiating a lesion (*V_max_*) is in the order of the single digit. These result echoes the number of virions transmitted by aphids (12, 13) and confirms the consequences of limited number of virions starting a new generation. In the case of the multiplication rate inside a lesion, the range is wider compared to the initiation of a lesion. This finding suggests that the host’s response, which includes various defense mechanisms, can vary in effectiveness and speed of activation.

**Fig. 5.**
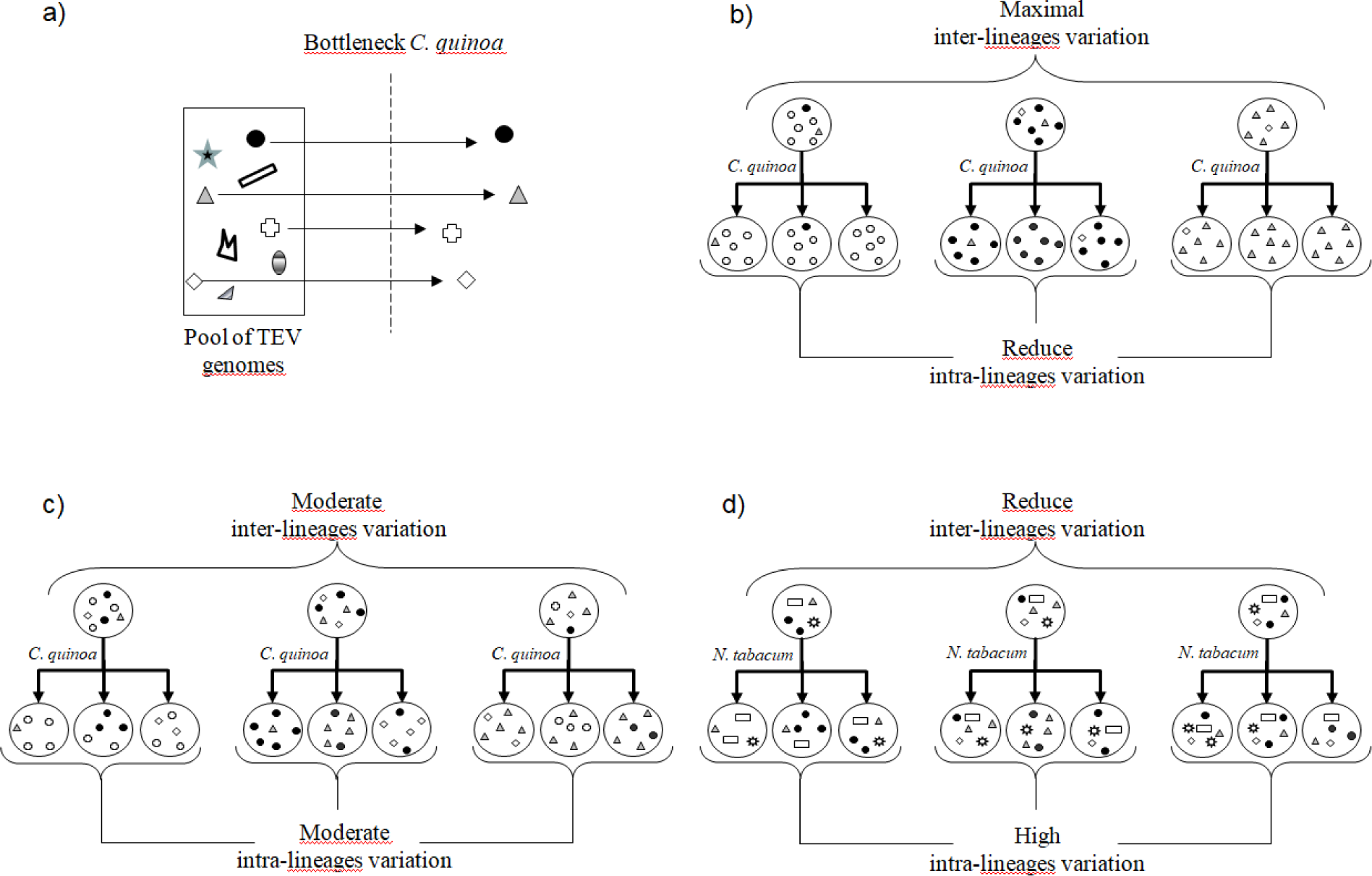
(A) Hypothetical effect of demographic bottlenecks of different size in the genetic diversity intra- and inter-linages after passage in *C. quinoa* after on passage. (B) For one lesion, (C) a group of lesions or (d) including a large population expansion in a systemic host such as *N. tabacum*.

Fig. 6 depicts model of the effect of bottlenecks in plant virus transmission. After a first bottleneck in *C. quinoa* (Fig. 6A) the inter-lineage genetic variability was different for each demographic condition. For the case of small *N_e_*, the founder effect resulted in a maximal inter-lineage variation (Fig. 6B), a moderate inter-lineage variation of infectivity for large *N_e_* dynamic (Fig. 6C) and a reduce inter-lineage variation for large *N_e_* dynamics together with a generation of diversity (Fig. 6D). The intra-lineage diversity produced is small, moderate and larger for small, intermediate and large *N_e_*demographics respectively.

In the light of these results, combination of a founder effect with a better multiplication in *C. quinoa* of one variant can explain the rate of infectivity evolution of the small and the intermediate *N_e_* modality, in line with the theoretical predictions by Bachtrog and Gordo (60).

Because of Muller’s ratchet effect, it can be anticipated that an asexual organism with high rate of mutation and small population size will not recover its initial fitness (22, 34, 58). Muller’s ratchet effects can be slowed or stopped by various factors as (*i*) increasing population size, (*ii*) epistatic selection and efficient purge of deleterious genomes (59), (*iii*) reducing the mutation rate, (*iv*) the appearance of compensatory or back-mutation (60, 62), and (*v*) recombination (59). In our experimental model, passaging of local lesions in *C. quinoa* results in a drastic reduction of the probability that such events will occur. Indeed, the virus population is contained in few cells and cannot expand. Given the small population size low generation number, the distribution of mutational (49, 50) and recombination effects (63), the probability that any fortuitous genetic changes be beneficial and not neutral or deleterious is very low.

RNA viruses encounter different bottlenecks during infection and expansion in plants, from inoculation by vectors to cell-cell and systemic movement (16). In this work, by modulating the bottleneck size through single/grouped lesions and multiplication in *N. tabacum*, we have been able to mimic the variety of bottlenecks encountered by the virus during its natural cycle. These bottlenecks may allow the virus to explore the fitness landscape by sampling viral population in the plant (7). Moreover, in case of cellular co-infection, fitness loss during repeated narrow genetic bottlenecks is an essential element of rapid selection. Plant RNA viruses rather than suffering from fitness losses use this mechanism to favor their adaptive variants (7, 8, 10). Moreover, when populations regularly experience bottlenecks, such as viruses, increasing the fitness cost of mutations so that unfit mutants are less likely to fix at each passage is an adaptative solution (64, 65). This approach to quickly remove non-viable genomes is affected when serial bottlenecks are experimented and moreover in a new environment leading to a weaker selection. In this study, we show that viral populations draw their strength from their ability to generate large populations. This ability coupled with high fitness costs during mutations allows facing the various bottlenecks encountered in a host (7).

It has been demonstrated that the average fitness of subclones does not predict the fitness of the whole clonal population at early time point (38, 54). This result underlines the advantage of grouping several lesions to avoid fitness losses. Moreover, this study has shown that differences in fitness between subclones and the population disappear after a long period of replication; in our case evolution of populations’ fitness in *C. quinoa* is stopped, although in *N. tabacum* the fitness can evolve.

These results highlight the combined effects of transmission bottlenecks, multiplication rate and selection in virus evolution in a plant system, as well as the life history of the founders. This mutation-selection balance can maximize adaptability while maintaining genetic structure of the virus population (65). Our data reinforce theoretical studies that conclude the mutation rate evolved in concert with the number of replication rounds coupled to bottlenecks events, in our case in *N. tabacum*. Therefore, a single founder particle is able to generate an expanding virus population.

In conclusion, we provide experimental and computational support to the hypothesis that exponential growth following strong population bottlenecks can stop the clicking of Muller’s ratchet, preventing the viral fitness declines. Such population expansions not only allow for the appearance and rise in frequency of compensatory and back mutations but also enhance the effectiveness of natural selection to fish out new beneficial mutations.

## ACKNOWLEDGEMENTS

We thank Francisca de la Iglesia for excellent technical assistance. SFE was supported by grants PID2022-136912NB-I00 funded by MCIN/AEI/10.13039/501100011033 and “ERDF a way of making Europe” and by CIPROM/2022/59 funded by Generalitat Valenciana. We are grateful to the genotoul bioinformatics platform Toulouse Occitanie (Bioinfo Genotoul, https://doi.org/10.15454/1.5572369328961167E12) and the computing facilities MCIA (Mésocentre de Calcul Intensif Aquitain) of the Université de Bordeaux and of the Université de Pau et des Pays de l’Adour for providing help and computing resources.

